# *Euprymna berryi* as a comparative model host for *Vibrio fischeri* light organ symbiosis

**DOI:** 10.1101/2025.01.10.632448

**Authors:** Avery M. Imes, Morgan N. Pavelsky, Klodia Badal, Derrick L. Kamp, John L. Briseño, Taylor Sakmar, Miranda A. Vogt, Spencer V. Nyholm, Elizabeth A. C. Heath-Heckman, Bret Grasse, Alecia N. Septer, Mark J. Mandel

**Affiliations:** Department of Medical Microbiology and Immunology, University of Wisconsin-Madison, Madison, WI USA; Genetics Training Program, University of Wisconsin-Madison, Madison, WI USA; Department of Earth, Marine & Environmental Sciences, University of North Carolina, Chapel Hill, NC USA; Department of Molecular and Cell Biology, University of Connecticut, Storrs, CT USA; Eugene Bell Center, Marine Biological Laboratory, Woods Hole, MA USA; Department of Integrative Biology, Michigan State University, East Lansing, MI USA

**Keywords:** bobtail squid, *Aliivibrio fischeri*, biofilm, motility, Type 6 Secretion System, apoptosis, laboratory models

## Abstract

Functional studies of host-microbe interactions benefit from natural model systems that enable exploration of molecular mechanisms at the host-microbe interface. Bioluminescent *Vibrio fischeri* colonize the light organ of the Hawaiian bobtail squid, *Euprymna scolopes*, and this binary model has enabled advances in understanding host-microbe communication, colonization specificity, *in vivo* biofilms, intraspecific competition, and quorum sensing. The hummingbird bobtail squid, *Euprymna berryi,* can be generationally bred and maintained in lab settings and has had multiple genes deleted by CRISPR approaches. The prospect of expanding the utility of the light organ model system by producing multigenerational host lines led us to determine the extent to which the *E. berryi* light organ symbiosis parallels known processes in *E. scolopes*. However, the nature of the *E. berryi* light organ, including its microbial constituency and specificity for microbial partners, have not been examined. In this report, we isolate bacteria from *E. berryi* animals and tank water. Assays of bacterial behaviors required in the host, as well as host responses to bacterial colonization, illustrate largely parallel phenotypes in *E. berryi* and *E. scolopes* hatchlings. This study reveals *E. berryi* to be a valuable comparative model to complement studies in *E. scolopes*.

**IMPORTANCE:** Microbiome studies have been substantially advanced by model systems that enable functional interrogation of the roles of the partners and the molecular communication between those partners. The *Euprymna scolopes-Vibrio fischeri* system has contributed foundational knowledge, revealing key roles for bacterial quorum sensing broadly and in animal hosts, for bacteria in stimulating animal development, for bacterial motility in accessing host sites, and for *in vivo* biofilm formation in development and specificity of an animal’s microbiome. *Euprymna berryi* is a second bobtail squid host, and one that has recently been shown to be robust to laboratory husbandry and amenable to gene knockout. This study identifies *E. berryi* as a strong symbiosis model host due to features that are conserved with those of *E. scolopes*, which will enable extension of functional studies in bobtail squid symbioses.

## INTRODUCTION

Symbiotic microbes play critical roles in animal development, immunity, and health. Invertebrate animal models have played foundational roles in elucidating the mechanisms by which microbiomes are established and maintained, and the bobtail squid-*Vibrio fischeri* system has been extensively studied with regard to host, symbiont, and the interaction between the two (1–5). Many bobtail squid species harbor a light-emitting organ that is colonized by luminescent bacteria, the light from which camouflages the host’s silhouette as the animal forages under moonlight (6, 7). During each generation, squid hatch aposymbiotically and acquire their symbiotic bacteria from the seawater. Over the course of hours, the colonization initiation process selects for the partner symbionts. Host-microbe communication continues into the persistence phase as the host provides nutrients to the bacteria, which in turn produce light for the host. A majority of symbionts are expelled each day and the remaining bacteria continue on a diel cycle of regrowth and luminescence (8, 9).

Most bobtail squid studies have focused on the Hawaiian bobtail squid, *Euprymna scolopes*. Studies in *E. scolopes* have revealed bacterial factors that are required for colonization initiation (10, 11), including flagellar motility (12, 13) and symbiotic biofilm formation (14–16). Luminescence, which is produced at high cell density in response to quorum sensing, is required during the persistence phase; while dark mutants can colonize successfully at 24 h, beginning at 48 h they exhibit a defect and are not detectable after a few weeks (17–19). Work in *E. scolopes* has additionally revealed principles regarding host responses to bacterial colonization. Ciliated epithelial appendages that recruit the symbionts undergo apoptosis and regression in response to release of the bacterial peptidoglycan fragment tracheal cytotoxin (TCT), with apoptotic nuclei detectable as early as 24 h post infection (20–22). Additionally, TCT triggers the migration of macrophage-like hemocytes into the light organ, where the hemocytes begin a light organ maturation process that results in the stable tolerance of the *V. fischeri* symbiont (23–25). As a result of the cited studies and many others, it is clear that there is a rich dialogue between host and microbe during the colonization process (26).

The availability of multiple bacterial strains has facilitated comparative approaches that have yielded a broad understanding of microbiome dynamics and evolution. Interbacterial competition takes place inside the host, as revealed from the function of a Type 6 Secretion System on chromosome II (T6SS2) that is present in approximately half of sequenced *V. fischeri* strains and enables those strains to kill competitors that lack the system (27, 28). Comparisons between fish and squid symbionts revealed differential biofilm regulation as a key basis for host colonization specificity in the North Pacific Ocean (29). At the same time, a variety of cephalopod models have been cultivated both within and outside the squid-Vibrio field; *E. scolopes* have been used to build cephalopod reference genomes (30, 31), multiple *Euprymna* species have revealed novel biology in neuroscience (32, 33), and a broad range of cephalopods have been used to study transcriptome plasticity (34). In contrast to the diversity of symbiont strains that are gaining traction for functional studies, most host work continues in *E. scolopes*. This is due to its substantial literature, ease of collection in the United States, and that it lays large clutches of eggs to support experimentation. While significant work has been conducted in other bobtail squid hosts, most notably *Euprymna tasmanica* (though also *Sepiola robusta* and others), no comparative host has offered scientific or husbandry advantages that have led to widespread adoption (35–37).

*Euprymna berryi*, the hummingbird bobtail squid, has been suggested as a cultivable model that could be applied for symbiosis studies (38). Recent efforts have expanded the aquaculture viability of valuable cephalopod models including *E. berryi* (38). Previous work culturing *E. berryi* noted similarity between *E. berryi* and *E. scolopes* and suggested that *E. berryi* could make a promising model system due to higher survivorship and fecundity in aquaculture compared to *E. scolopes* (38). The first CRISPR-mediated knockouts in bobtail squids were performed in *E. berryi* to remove pigmentation from the organism, demonstrating the ability to genetically manipulate *E. berryi* (39). Therefore, the focus of this study is to evaluate the suitability of lab-raised *E. berryi* for mechanistic studies in an animal-microbe symbiosis.

## RESULTS

### *Euprymna berryi* bobtail squid harbor luminescent bacteria in a light organ

Given the prominence of *E. scolopes* as a model host for light organ symbioses, we directly compared the adult and hatchling *E. berryi* animals with their *E. scolopes* counterparts. While similar in overall appearance, the *E. berryi* animals were larger than the *E. scolopes* animals (**Fig. 1, Fig. S1**), and the *E. berryi* animals were darker brown in color compared to *E. scolopes* adults when the animals were placed on the same substrate. The eggs laid by *E. berryi* animals were also darker and had more defined structures outside of the eggs. However, the hatchling animals of both species were similar in color, with the *E. berryi* hatchlings notably larger (**Fig. 1**).

**Figure 1.**
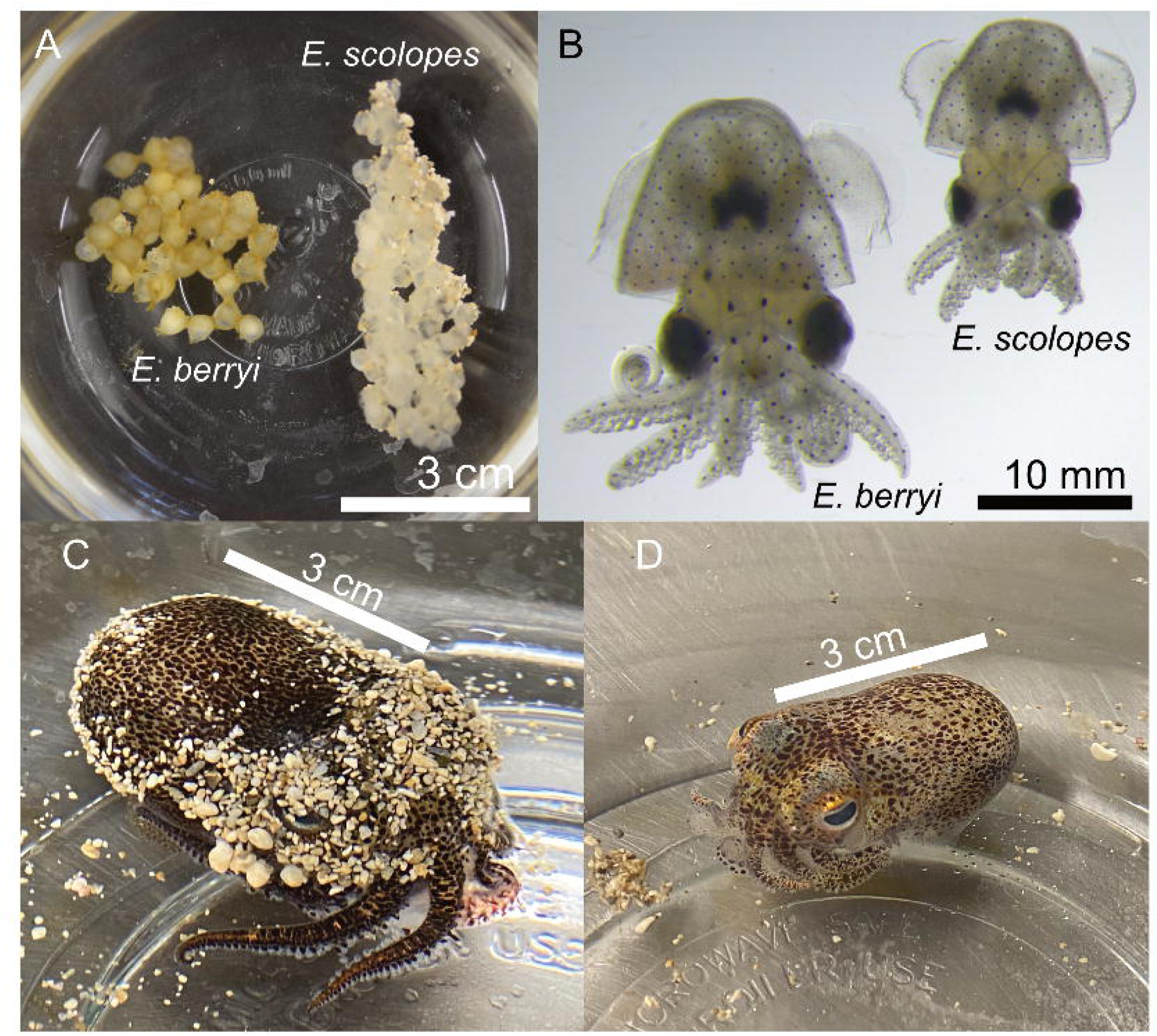
Comparison of *Euprymna* species. (A) *E. berryi* (left) and *E. scolopes* (right) eggs immediately prior to hatching. (B) An *E. berryi* and *E. scolopes* hatchling 24 hours post-hatching. (C) An adult female *E. berryi.* (D) An adult female *E. scolopes*.

We used GFP-expressing *V. fischeri* strains as biological probes to detect a possible light organ in *E. berryi.* The strains used included the *E. scolopes* light organ isolate ES114 and the *E. berryi* isolate BW2 that we describe in detail later. *E. berryi* were reported to contain a luminous organ when first identified (40), so we asked whether this organ was colonized by bioluminescent bacteria as observed in closely related *Euprymna* species. We introduced the bacteria to water containing aposymbiotic *E. berryi* hatchlings, washed the hatchlings at 3 and 24 h, and assayed luminescence of the animals at 48 h. All animals were luminous and were fixed for imaging. Confocal imaging of the animals revealed that the bacteria of both strains were localized to the light organ in the mantle cavity in a structure that strongly resembled the light organ of *E. scolopes* (41). Representative images from BW2 are presented in Fig. 2 and quantification of the distribution of both strains are presented in Fig. S2. The bilobed organ of *E. berryi* contains three crypts on each side, with a dedicated exterior pore connecting to each crypt. The posterior pore leads to Crypt 2 (c2), middle pore to c1, and anterior pore to c3 (Fig. 2), which matches the morphology observed in *E. scolopes* (41). The relative crypt size is conserved with *E. scolopes*, with c1 as the dominant crypt, and c2 as the next smallest followed by c3 (Fig. S2). c1 contains approximately 70% of the bacteria by volume (Fig. S2). Furthermore, we observed antechambers and host anatomical bottleneck structures that were first described in *E. scolopes* (Fig. 2C) (41). Overall, imaging suggested strong conservation of light organ structures between *E. scolopes* and *E. berryi*.

**Figure 2.**
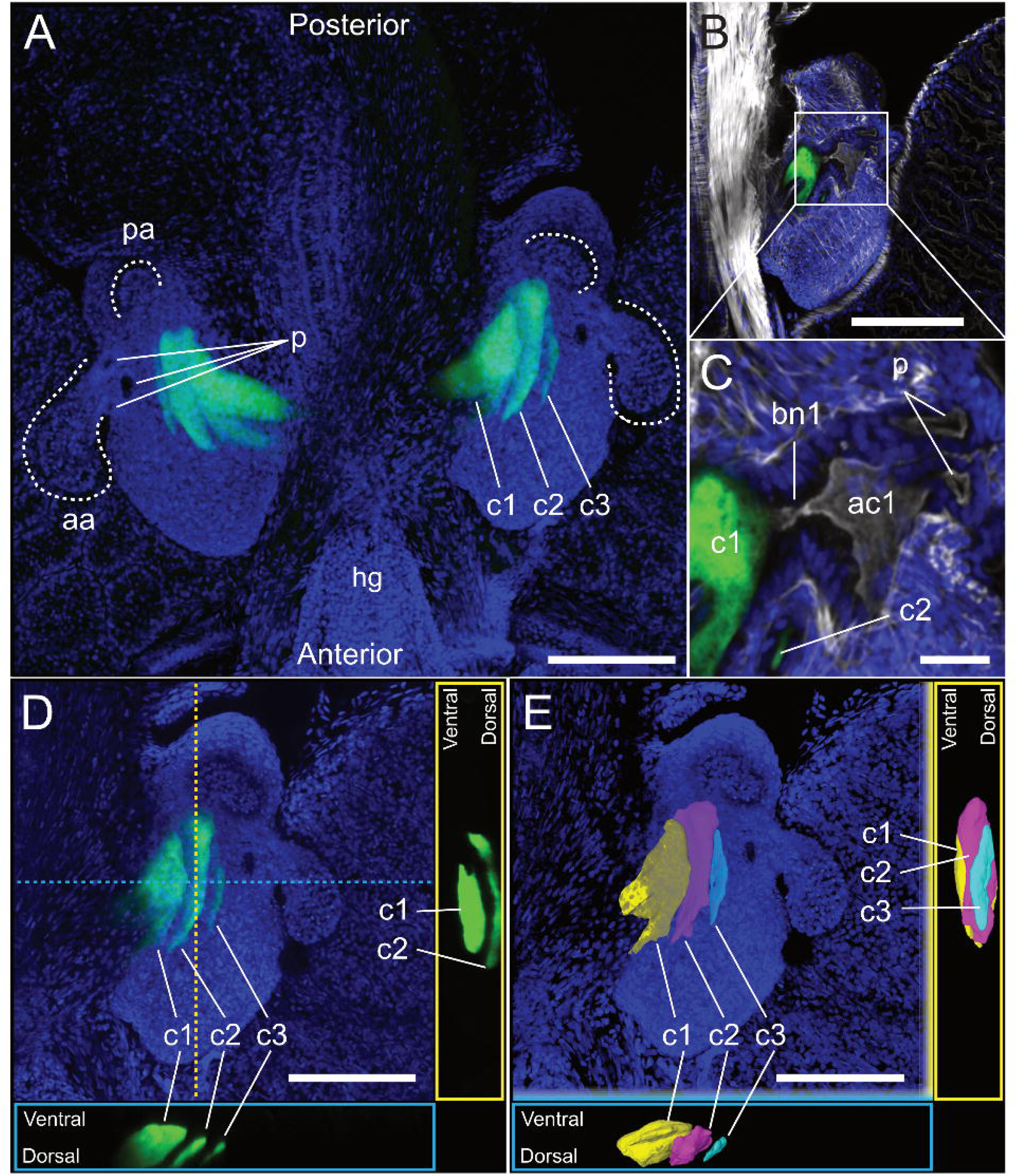
The light organ of juvenile *E. berryi*. Images are representative of all symbiotic animals (*n* = 10). (A) Representative image of an *E. berryi* light organ. Image is a maximum intensity projection of a confocal z-stack approximately 150 µm deep. DAPI staining (blue) of host tissue shows that each side of the bi-lobed organ has a long anterior appendage (aa) and shorter posterior appendage (pa). At the base of each anterior appendage are three pores (p). Three crypts (c1-3) are on each side. Each crypt is visualized by GFP-expressing *V. fischeri* (green). The hindgut (hg) lies in the anterior end. (B) A single lobe of the light organ, counterstained with Alexa640-conjugated phalloidin to visualize actin cytoskeleton. (C) Enlarged image of boxed region in B. Pores (p) lead to crypts. A tight bottleneck (bn) separates the antechamber (ac) from crypt 1 (c1). (D) Maximum intensity projection and orthogonal views of crypts from one lobe of the light organ. Locations of orthogonal slices represented by dashed line. Each of the three crypts (c1, c2, c3) is fully colonized by GFP-expressing *V. fischeri*. (E) Three-dimensional model of the crypts, each a different color. Crypt 1 (c1) is the largest crypt, with branching structures. The smaller crypts 2 (c2) and 3 (c3) lay dorsally to crypt 1. Bars: A&B = 150 µm, C = 10 µm, D&E = 100 µm.

### *V. fischeri* and other γ-proteobacteria were isolated from *E. berryi* adult tank water and colonized hatchlings

We sought to identify bacterial strains that colonize the *E. berryi* light organ. We noticed that *E. berryi* raised with access to water pumped in from Eel Pond (Woods Hole, MA, USA) readily became luminous. Therefore, we used bacteria isolated from these luminous *E. berryi* hatchlings as a source for putative *E. berryi* colonizing strains. As a second source for bacterial isolates, we hypothesized that *E. berryi* may expel bacteria in a rhythmic daily cycle similar to *E. scolopes* (9, 23, 42), so we isolated bacteria from the tank water of wild-caught *E. berryi* adult animals. In both cases, we isolated bacteria on LBS agar, streak-purified different colony morphologies, and conducted secondary screening for isolates that were detected as luminous in liquid LBS medium. We identified nine (9) distinct strains from the adult “*berryi* water” (BW) sampling: two of these isolates represented colony types that were highly abundant in our sampling yet minimally luminous (BW4, BW8), while the other seven were luminous. From the “*berryi* hatchling” (BH) sampling, we identified two (2) strains, both of which were luminous. For each strain above, we grew a culture in LBS media, obtained whole genome Nanopore sequencing, and determined the likely species by FastANI analysis (43). This analysis identified three *V. fischeri* strains (BH2, BW2, BW9), 7 other *Vibrio spp.* isolates, and one *Pseudoalteromonas flavipulchra*. Figure 3 lists the isolates and shows specific luminescence for each strain when grown in liquid culture.

**Figure 3.**
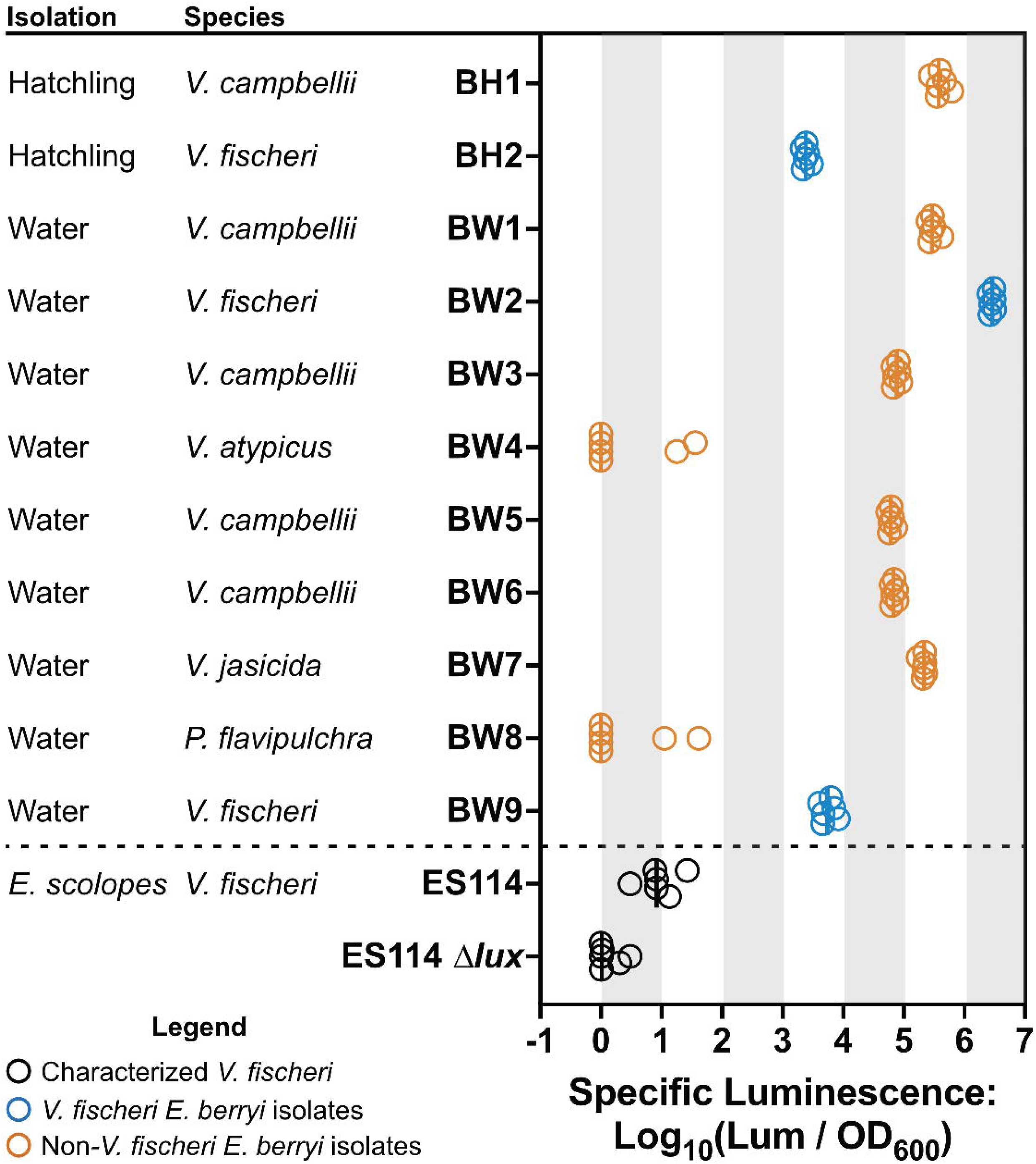
Isolation and characterization of bacterial strains. Hatchling isolates were plated from colonized *E. berryi* hatchlings. Water isolates were plated from tank water housing a wild-caught adult *E. berryi*. Isolate species was identified by average nucleotide identity. Specific luminescence for each strain (*n* = 6) when grown in liquid culture was plotted with circles representing individual biological replicates. The median is indicated by a vertical bar.

### *E. berryi* is colonized specifically by *V. fischeri* bacteria

The *E. scolopes* light organ is colonized only by *V. fischeri* bacteria, yet we isolated a number of bacterial species from the water and hatchlings. We therefore asked whether *E. berryi* was more permissive to colonization. We conducted standard 3 h inoculation, 48 h colonization assays that are well-established for *E. scolopes* and that we adapted for *E. berryi* (44). The key adjustments needed from our published protocol were to accommodate the larger animal size. Specifically, we enabled a larger opening into the transfer pipet by using a razor blade to cut the stem of the transfer pipet closer to the bulb, where the stem diameter is larger, and we transferred the animals at a slower pace, which we found necessary with the larger bodies.

We found that our *E. berryi*-isolated *V. fischeri* strains generally colonized *E. berryi* hatchlings, with BW9 as the only strain showing slightly reduced CFU counts (Fig. 4A). In contrast, the other species isolated from *E. berryi* water were unable to colonize, as was the *V. campbellii* isolate BH1 isolated from a hatchling. These colonization patterns match what has been reported for colonization of non-*V. fischeri* strains in *E. scolopes* (45, 46).

**Figure 4.**
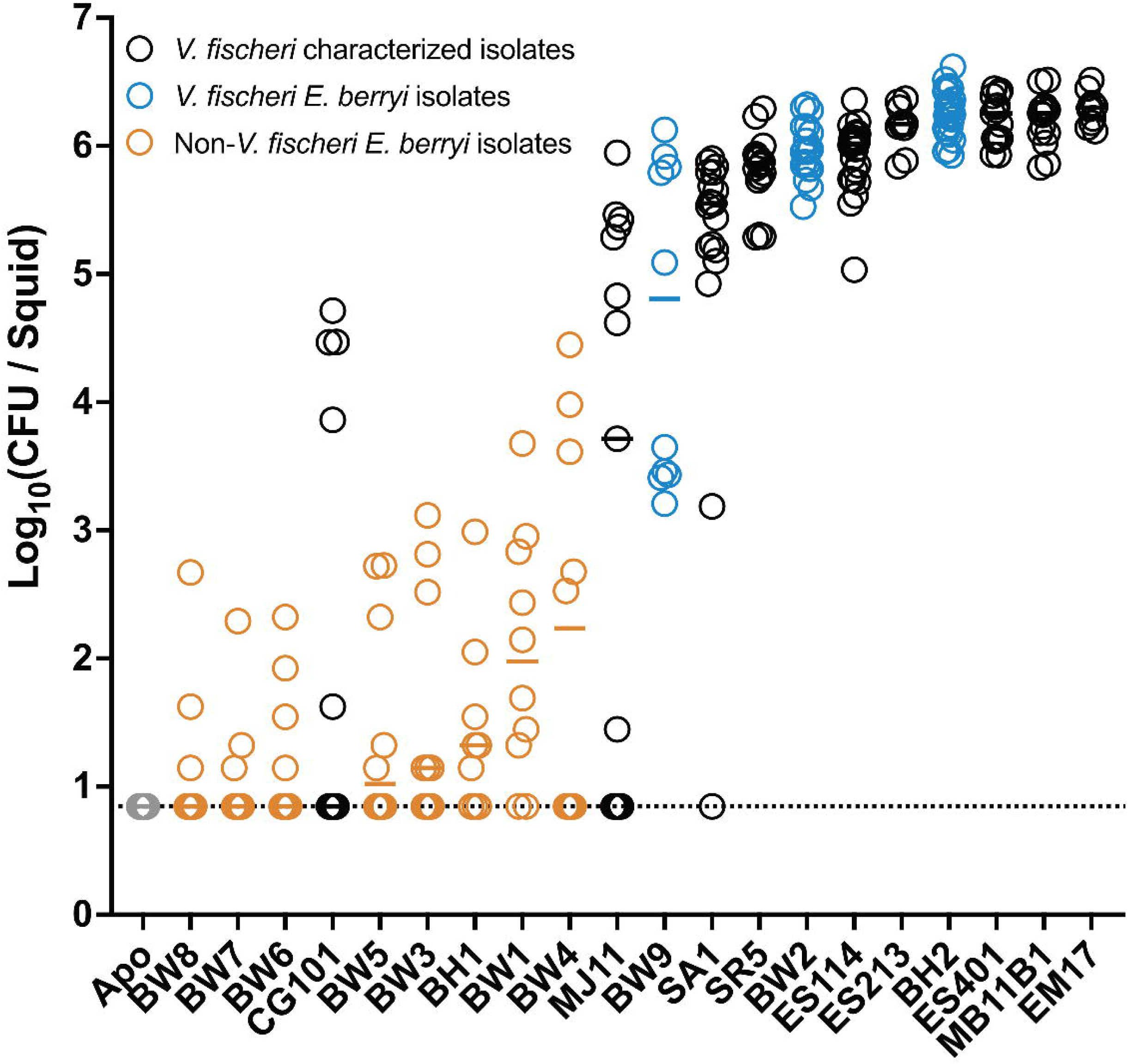
*V. fischeri* are the exclusive colonizers of *E. berryi*. *E. berryi* hatchlings were used in single-strain colonization experiments, where circles represent individual animals. The limit of detection for this assay, represented by the dashed line, is 7 CFU/light organ. Horizontal bars represent the median for each group (*n* = 10-25).

Our standard method of assaying colonization includes homogenization of thawed frozen whole animals, as symbionts in the light organ are osmotically protected from freezing while other bacteria in the animal are generally at low abundance and do not survive freezing well. However, to gain additional precision into the patterns observed in Figure 4, we dissected light organs from squid inoculated with a subset of the strains. Those data, shown in Fig. S3A, demonstrate robust light organ colonization for the new *V. fischeri* strains, with the same relative abundance of symbionts across the three samples. Non-*V. fischeri* strains BW1 and BW4 show essentially no light organ colonization. Fish *V. fischeri* isolates CG101 and MJ11 demonstrate either no colonization or low colonization, consistent with their data from the whole animal assay. Additionally, we inoculated hatchlings with GFP-expressing derivatives of the same strains to visualize bacterial localization. We observed consistent presence of the new *V. fischeri* strains BH2, BW2 and BW9 in the light organ, but did not detect CG101, MJ11, BW1 or BW4 in the light organ (Fig. S3B). Overall, these data support a clear specificity for *V. fischeri*, and squid isolates of *V. fischeri*, in colonization of the *E. berryi* light organ.

*V. fischeri* isolated from squid light organs in the North Pacific Ocean and the Mediterranean Sea that can robustly colonize *E. scolopes* are similarly able to colonize *E. berryi* (Fig. 4A). This includes *V. fischeri* from *E. scolopes, Euprymna morsei*, *Sepiola robusta, and Sepiola affinis*.

The only divergence between colonization patterns in *E. scolopes* and those in *E. berryi* were noted for the fish isolates MJ11 and CG101. These strains do not robustly colonize the Hawaiian host (29, 47, 48), yet for both strains we observed a bimodal colonization pattern of *E. berryi* (Fig. 4A). In both cases, most animals had either undetectable levels of bacteria or >10^3^ CFU per squid. While in the case of MJ11, the bimodality partitioned by experiment, in neither case did it correlate with the concentration of bacteria in the inoculum (Fig. 4A, Fig. S4).

The above results suggest that most colonization patterns for *E. scolopes* strains are conserved during colonization in *E. berryi*. We therefore asked whether the new *V. fischeri* isolates from *E. berryi* were capable of colonizing *E. scolopes* squid. Not all *V. fischeri* can colonize *E. scolopes*, although squid isolates of *V. fischeri* (including those from other bobtail squid species) are typically able to colonize *E. scolopes* (29, 47, 49, 50). All three of the new *V. fischeri* strains isolated from *E. berryi* samples, BW2, BW9, and BH2, consistently colonized *E. scolopes* squid (Fig. S5).

### Bacterial factors required for host colonization in *E. scolopes* are similarly required for colonization of *E. berryi*

Decades of work in *E. scolopes* have defined bacterial colonization factors (10, 11, 13–15, 51–53). If the process of colonization is conserved in *E. berryi*, then a specific prediction is that mutants that fail to undergo key steps in the initiation or persistence of the symbiosis will exhibit colonization defects in *E. berryi*. To test this hypothesis, we examined ES114 mutants with initiation defects in the Syp biofilm (Δ*sypF*) or in flagellar motility (Δ*flrA*) and determined their ability to colonize *E. berryi*. Both genes are required for colonization of *E. scolopes* (13, 15, 51, 53). Both strains exhibited significant colonization defects with a majority of animals completely uncolonized (Fig. 5). In *E. scolopes*, non-luminescent mutants such as Δ*luxCDABEG* (Δ*lux*) are known to colonize the light organ but fail to persist. We observed a similar defect in *E. berryi*, with no significant defect at 24 h post inoculation (hpi), but with a highly significant defect and 4-fold fewer bacteria at 48 hpi (Fig. 5). This pattern matches that observed in *E. scolopes,* wherein the Δ*lux* mutant first exhibits its persistence defect at 48 hpi and is eventually lost from the *E. scolopes* light organ over 3-4 weeks (17–19). These data together argue that colonization in *E. berryi* bears large parallels to that which occurs in *E. scolopes*, especially with regard to specificity for bacterial strains and genetic requirements during symbiont entry and establishment in the host.

**Figure 5.**
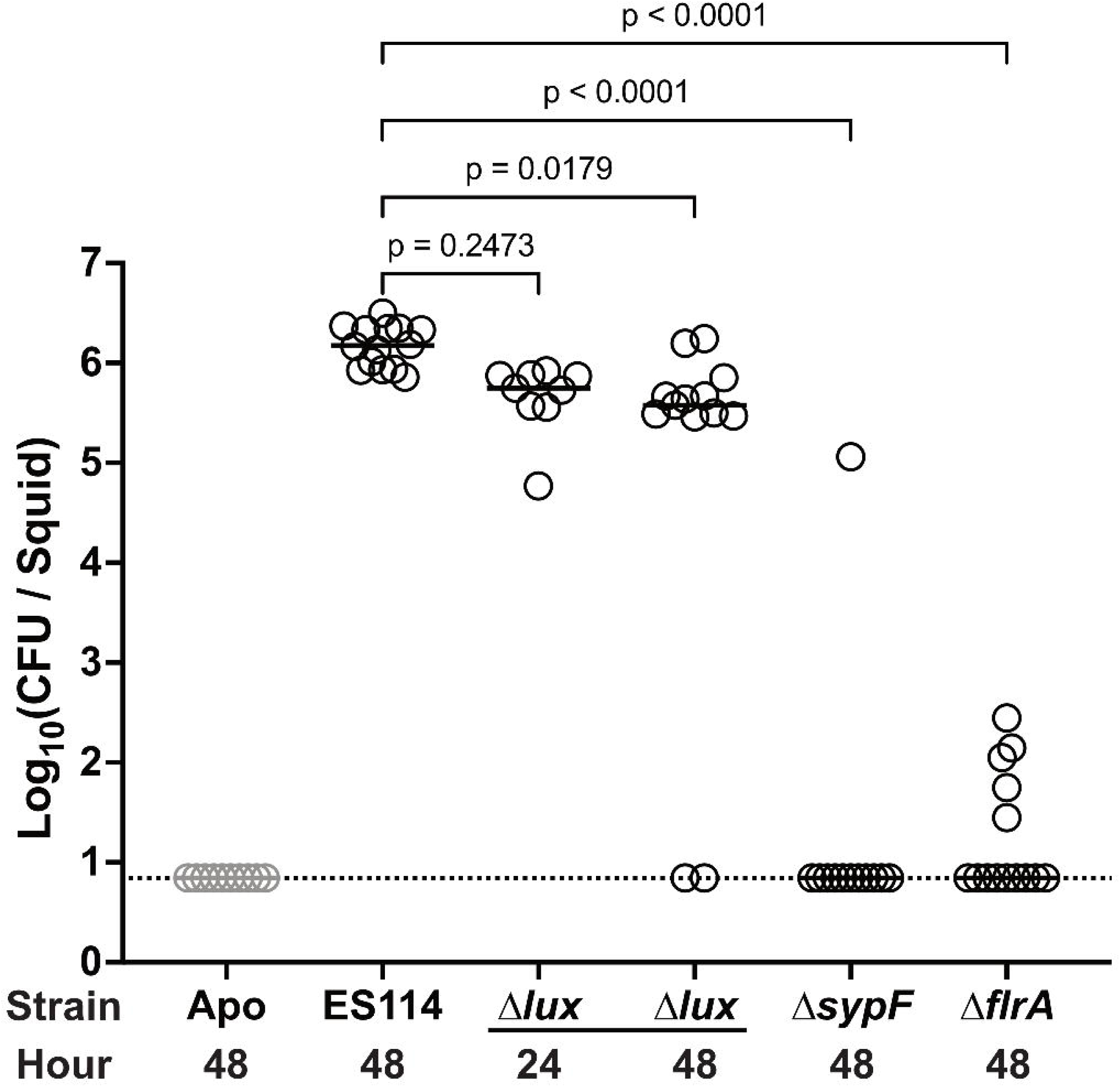
Colonization-deficient mutants are defective in *E. berryi*. *E. berryi* hatchlings were used in single-strain colonization experiments, where circles represent individual animals. The limit of detection for this assay, represented by the dashed line, is 7 CFU/light organ. Horizontal bars represent the median for each group. All mutants are constructed from the wildtype strain ES114. The inoculum strain is indicated on the X-axis, with the hour squid were fixed post-inoculation labeled below each group (*n* = 9-13). Treatment groups were compared to the ES114 control group using a Kruskal-Wallis test using Dunn’s multiple comparisons test.

### *The E. berryi* appendages exhibit developmental phenotypes cued by bacterial colonization

In *E. scolopes*, colonization by the *V. fischeri* symbiont triggers irreversible host morphological changes, particularly the dramatic apoptosis of the light organ appendages (21). In *E. scolopes,* colonized squid have detectable TUNEL staining in the anterior appendage at 24 hpi as a result of DNA fragmentation in apoptotic cells. To determine if similar host development is triggered by symbiotic colonization in *E. berryi*, we used a TUNEL assay. In *E. berryi,* BW9 induced the highest TUNEL count, followed by BH2 (Fig. 6A). Another measure for host development stimulated by symbiont colonization is hemocyte trafficking into the light organ appendages, which is elevated at 24 hpi in *E. scolopes* and minimal in aposymbiotic animals. Among the strains tested, BW9 and BH2 again promoted the strongest responses but with a wide range of variability across animals in all groups (Fig. 6B). BW2 was the only strain that did not yield statistically significant hemocyte trafficking (Fig. 6B). Overall, these results point to conserved host responses in *E. berryi* compared to *E. scolopes*, and with BH2 and BW9 (along with ES114) able to elicit both TUNEL staining and hemocyte trafficking in *E. berryi*.

**Figure 6.**
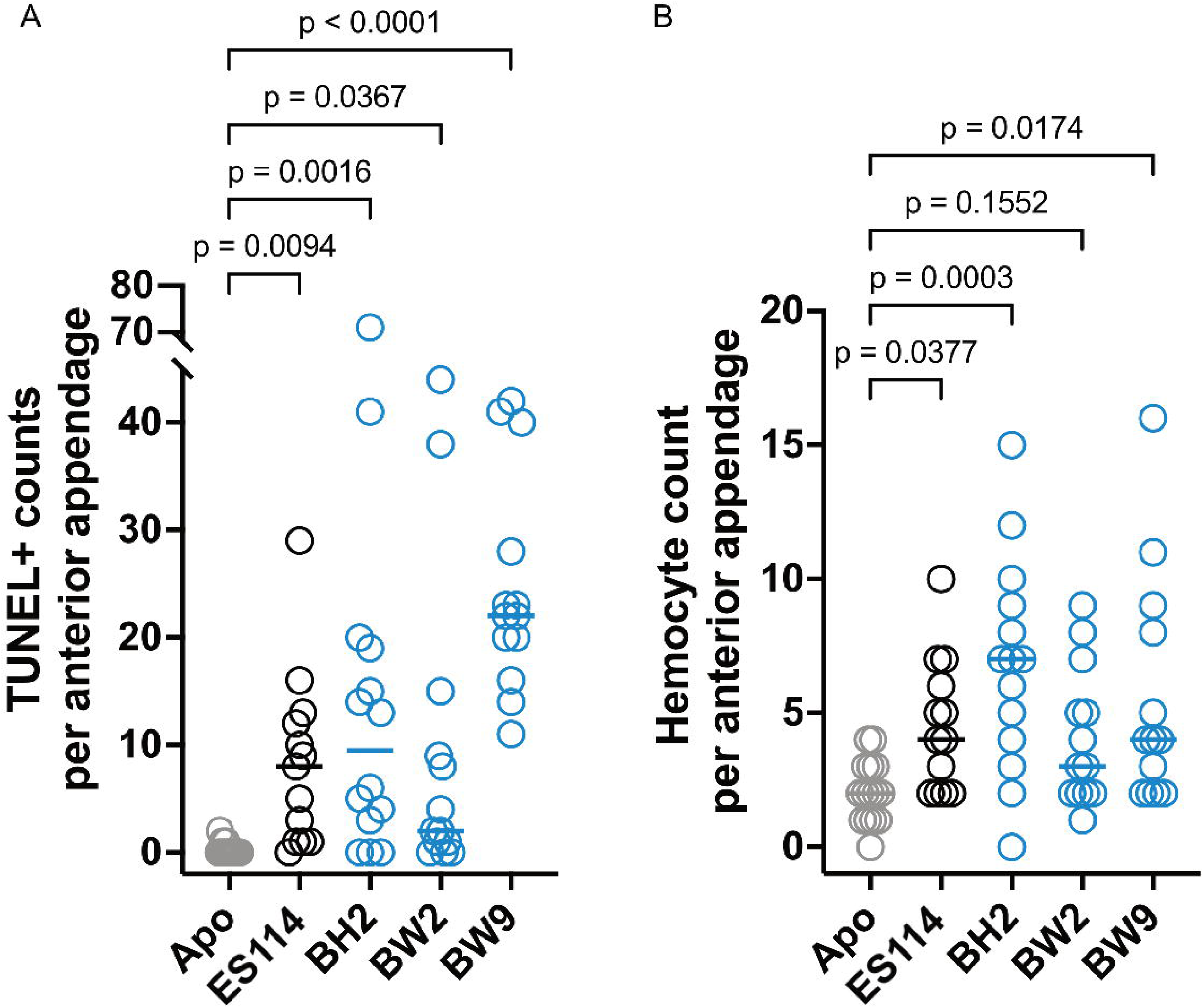
*V. fischeri* strains stimulate variable intensities of host development responses in *E. berryi*. *E. berryi* hatchlings were inoculated with a single bacterial strain and evaluated at 24 hours post-inoculation, where circles represent individual animals. Horizontal bars represent the median for each group (*n* = 13-15). New bacterial isolates from *E. berryi* are shown as blue circles. (A) The TUNEL+ counts in the anterior appendage were taken as a measurement for cell apoptosis, a marker of colonization-driven host development. (B) Hemocyte counts in the anterior appendage were used to measure hemocyte trafficking into the appendage, which is elevated in colonized *E. scolopes*. P-values were calculated using a Kruskal-Wallis test with Dunn’s multiple comparisons test.

### Type 6 secretion-mediated competition operates in the *E. berryi* host

Among the competitive mechanisms influencing squid colonization, some *V. fischeri* strains are known to use a strain-specific type VI secretion system found on chromosome II (T6SS2), which confers a contact-dependent ability to kill off competitive strains (27). We aimed to determine if this mechanism for interbacterial competition operates and influences colonization of *E. berryi*. All three *V. fischeri* isolates from *E. berryi* encode T6SS2. To determine if the system was functional, we co-incubated the isolates with GFP-expressing ES114. The control strain for this assay, the *E. scolopes* isolate ES401, kills the target ES114-GFP and results in no detectable fluorescence from the resulting colony spot. In the absence of the structural protein TssF, T6SS2 activity is prevented and ES114-GFP is able to grow and produce fluorescence. This phenotype was mirrored in the isolates from *E. berryi*: all three strains exhibited functional T6SS2 activity, and this activity was removed in BW2 by mutation of *tssF* (Fig. 7A). A halo effect seen in BW2—i.e., the green ring around the edge of the colony spot—has been previously observed in other T6SS2^+^ strains and sometimes occurs when survivors at the edge of the colony escape T6SS-mediated killing. Moreover, when BW2-derived strains expressed a GFP-tagged component of the sheath protein (VipA) that is required for assembly of the T6SS injection apparatus, sheaths were observed in BW2 as puncta near the edges of cells (Fig. 7B). However, for the *tssF* mutant that lacks the ability to make T6SS2 sheaths, sheaths were not observed, with the VipA-GFP protein instead forming a homogeneous distribution of the GFP across the cell (Fig. 7B) and results in a non-killing phenotype (Fig. 7AB).

**Figure 7.**
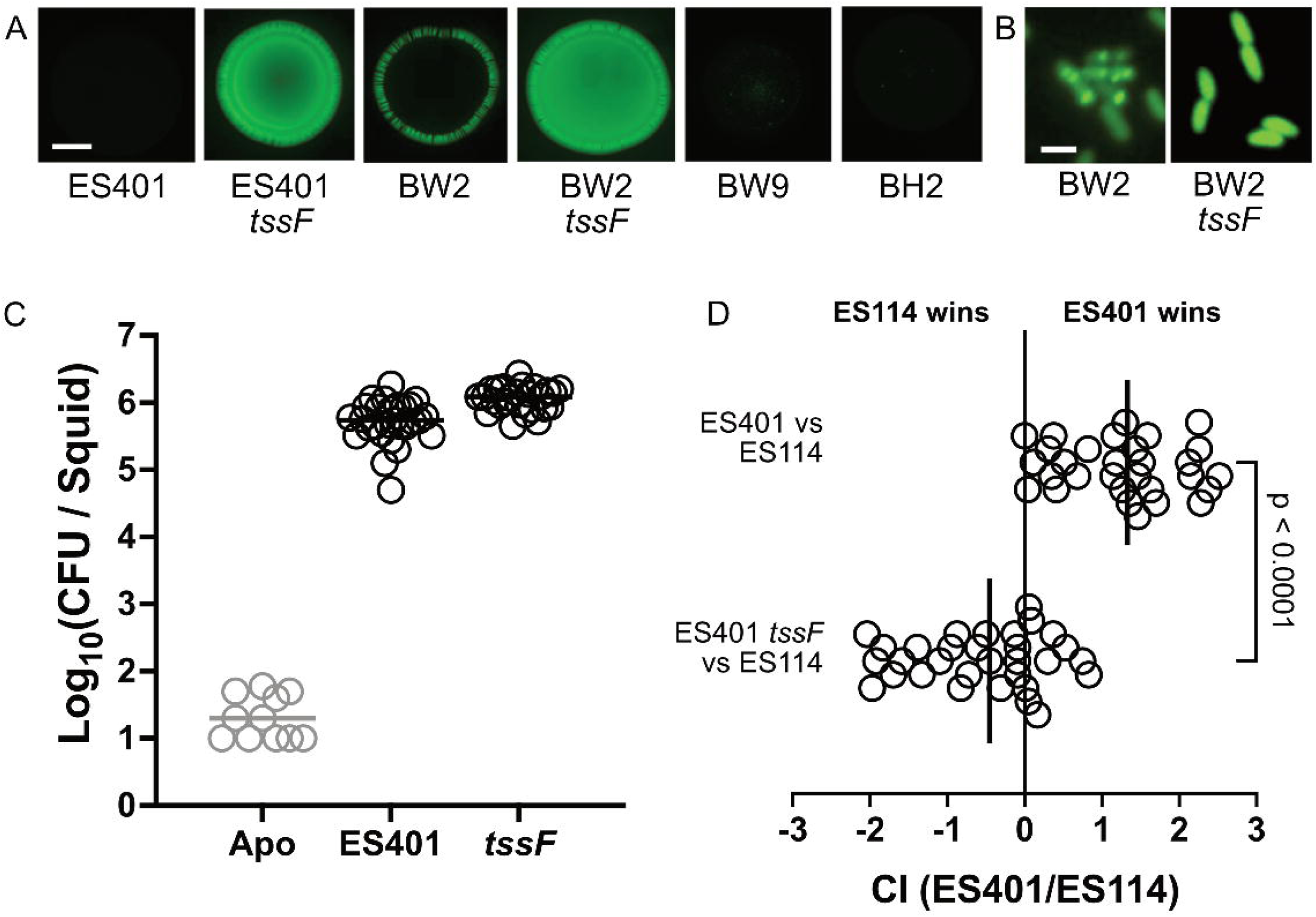
*E. berryi* can be used to study T6SS-mediated competition in *V. fischeri* symbionts. (A) Coincubation assays between the indicated strain and a GFP-tagged ES114 target strain. The scale bar for images in panel A is 2 mm. (B) Representative images of T6SS2 sheaths in wildtype BW2 and the *tssF* derivative carrying a VipA-GFP expression vector. The scale bar for images in panel B is 2 µm. (C) Single colonization assays for indicated ES401 strains at 24 hr (*n* = 11-28). (D) Competitive colonization assays with GFP-tagged ES114 and RFP-tagged ES401-derived strain (*n* = 30-31 per competition). Time points are 24hr post-inoculation. Panel D shows the competitive index (CI) in log 10 scale for each squid, where the ending strain ratio is divided by the beginning strain ratio. Statistical comparison was made using a Mann-Whitney test.

To ask whether T6SS-mediated competition occurs in *E. berryi* juveniles, as it does in *E. scolopes* (54), we conducted competition assays within the squid host. Both ES401 and its non-killing *tssF* derivative are able to consistently colonize *E. berryi* in single-strain colonizations (Fig. 7C). To determine if T6SS2 competition functions in the *E. berryi* host, we used a competitive colonization assay between ES401 and ES114, where strains are differentially fluorescently labeled and co-inoculated into squid hatchlings. ES401 exhibited a 60-fold competitive colonization advantage over ES114 (Fig. 7D; comparing the *tssF*^+^/*tssF*^−^ results). This result confirmed that T6SS2 in the *V. fischeri* isolates from *E. berryi* is functional and that the system provides a competitive advantage in colonization in *E. berryi*.

## DISCUSSION

Overall, this work establishes *E. berryi* as a culturable model for bobtail squid-*V. fischeri* symbiosis studies. We demonstrate that for the light organ symbiosis, the host anatomy, host-microbe specificity, bacterial gene requirements, interbacterial competition, and host responses to symbiosis broadly mirror those processes observed in the well-studied *E. scolopes* model. Recent work in wild-caught *E. berryi* characterized the microbial community in the female accessory nidamental gland (ANG), a reproductive organ in which bacteria provide key defensive roles including provisioning antifungal compounds into the jelly coat of the developing squid eggs (55–58). Therefore, the ability to use cultured animals for bobtail squid symbiosis studies, in a host species in which it was recently shown that gene knockouts could be conducted (39), opens additional resources to investigate how animal microbiomes are established across both the light organ and ANG microbiomes.

One area in which we observed a variable response in *E. berryi* was in the host responses to symbiotic colonization. Although all strains tested exhibited a significant response in the TUNEL assay, the overall response was weak compared to *E. scolopes*, which averages around 40 TUNEL+ cells per anterior appendage in colonized animals at the 24 hpi time point (22). Only strain BW9 exhibited median counts of >20 per appendage (Fig. 6A). Similarly, the average hemocyte count induced by ES114 has been previously observed to be higher in *E. scolopes* than we observed in Fig. 6B, averaging 15 per anterior appendage (59). One possibility is that the host responses in *E. berryi* are qualitatively similar but quantitatively distinct from *E. scolopes*. Another possibility is that the timeline for the response in *E. berryi* differs from that in *E. scolopes*. A future study could examine this question by comparing the responses side-by-side, and carrying out the timeline to 5-7 d. A general limitation of this study is that most work was examined only to the 48 h timepoint, so differences at later times are possible and were not examined in this initial investigation.

We expect that the non-*V. fischeri* strains isolated from the *E. berryi* water and hatchlings were not light organ symbionts, but instead were environmental samples. We did not conduct light organ dissections for the BH samples, so bacteria present in other tissues could have been isolated if they were osmotically protected during freezing. Given that the non-*V. fischeri* strains failed to colonize aposymbiotic *E. berryi* hatchlings when presented at >10^3^ CFU/ml in monoassociation suggests that they are not able to colonize the light organ. The *V. fischeri* isolates BH2, BW2, and BW9 are clearly capable of colonizing the bobtail squid light organs, both for *E. berryi* and *E. scolopes*, and with BW2 we confirm specific light organ colonization via direct imaging (Fig. 2). These *V. fischeri* isolates may have originated from the adult *E. berryi* in the aquaculture system or from the Atlantic Ocean water feeding the tank systems. Strain BH2 colonizes *E. berryi* consistently across replicates (Fig. 4) and also stimulates significant host responses in both assays tested (Fig. 6), suggesting that it may be a natural symbiont and/or share characteristics with natural *E. berryi* symbionts. When it is possible to isolate symbionts from wild *E. berryi* animals, it would be valuable to address the phylogenetic distribution and phenotypes of those symbionts.

In summary, we describe *E. berryi* as a model for *V. fischeri* colonization and light organ symbiosis. This work therefore will enable use of tools developed for *E. berryi* to be applied for symbiosis studies and will allow for comparative studies between bobtail squid hosts.

## MATERIALS AND METHODS

### Animal comparison

#### Eggs/embryos

*Euprymna* spp. eggs were imaged using a Nikon D810 digital camera at University of Wisconsin-Madison and measured using FIJI (60). *Euprymna* spp. hatchlings were imaged within 24 hours post-hatching using a Leica M60 stereomicroscope with Leica FireCam software. **Adults.** Adult *E. berryi* (120-178 d post-hatching) were raised at Woods Hole Marine Biological Laboratory and photographed using an iPad 9th generation (model number MK2K3LL/A). Adult *E. scolopes* were collected from Oahu, Hawaii, and housed in an aquaculture facility for 61 d. Artificial seawater was maintained at a salinity of 34-35 ppt by addition of Instant Ocean (Spectrum Brands, Blacksburg, VA) to RO water. Animals were photographed using a Nikon D810 digital camera. Mantle lengths were measured using FIJI (60). A non-parametric one-way ANOVA (Kruskal-Wallis) using Dunn’s multiple comparisons test was used to compare the mantle lengths between males and females of each species.

### Bacterial strains and media

Bacterial strains used in this study are listed in Table 1. All bacteria except for *E. coli* were grown in Lysogeny broth salt (LBS) medium at 25°C (25 g Difco LB broth [BD], 10 g NaCl, and 50 mL 1 M Tris buffer [pH 7.5] per liter). An *E. coli* strain, used as the conjugation donor for the GFP plasmid, was grown in Lysogeny broth (LB) medium at 37°C (25 g Difco LB broth [BD] per liter). When required, kanamycin was added to a final concentration of 100 μg/ml (*V. fischeri*) or 50 μg/ml (*E. coli*), and erythromycin to 5 μg/ml (*V. fischeri*). *E. coli* strains requiring erythromycin were grown in Brain-Heart Infusion (BHI) medium at 37°C (37 g BBL BHI powder [BD] per liter) with 150 μg/ml. Growth media was solidified to 1.5% agar as needed by adding 15 g Bacto agar (BD) per liter.

**Table 1.** List of strains used in this study. Strains are listed by name, and lab stock number is shown in parentheses.

| Strain | Description | Source |
| --- | --- | --- |
| MJ11 (MJM1059) | Natural isolate, *Monocentris japonica* light organ | (49) |
| ES114 (MJM1100) | Natural isolate, *E. scolopes* light organ | (42) |
| CG101 (MJM1115) | Natural isolate, *Cleidopus gloriamaris* light organ | (47) |
| ES213 (MJM1117) | Natural isolate, *E. scolopes* light organ | (66) |
| MB11B1 (MJM1130) | Natural isolate, *E. scolopes* light organ | (67) |
| EM17 (MJM1136) | Natural isolate, *E. morsei* light organ | (68) |
| ES114 $\Delta luxCDABEG$ (MJM1296) | Non-luminescent ES114 (MJM1100) | (18) |
| ES401 (MJM1318) | Natural isolate, *E. scolopes* juvenile light organ | (47) |
| ES114 $\Delta flrA::bar$ (MJM3782) | Non-motile ES114, barcoded mutant (MJM1100) | (69) |
| ES114 $\Delta sypF::bar$ (MJM3972) | Syp-deficient ES114, barcoded mutant (MJM1100) | (70) |
| SA1 (MJM3859) | Natural isolate, *Sepiola affinis* light organ | (50) |
| SR5 (MJM3878) | Natural isolate, *Sepiola robusta* light organ | (50) |
| MJM1100 / pVSV102 (MJM1107) | ES114-GFP (Kan<sup>R</sup>) | (11) |
| MJM1059 / pVSV102 (MJM1112) | MJ11-GFP (Kan<sup>R</sup>) | This study |
| MJM5770 / pVSV102 (MJM5780) | BH2-GFP (Kan<sup>R</sup>) | This study |
| MJM5772 / pVSV102 (MJM5781) | BW2-GFP (Kan<sup>R</sup>) | This study |
| MJM5779 / pVSV102 (MJM5782) | BW9-GFP (Kan<sup>R</sup>) | This study |
| MJM1115 / pVSV102 (MJM6330) | CG101-GFP (Kan<sup>R</sup>) | This study |
| MJM5771 / pVSV102 (MJM6331) | BW1-GFP (Kan<sup>R</sup>) | This study |
| MJM5774 / pVSV102 (MJM6332) | BW4-GFP (Kan<sup>R</sup>) | This study |
| BH1 (MJM5769) | Natural isolate (*Vibrio campbellii*), *E. berryi* hatchling | This study |
| BH2 (MJM5770) | Natural isolate (*Vibrio fischeri*), *E. berryi* hatchling | This study |
| BW1 (MJM5771) | Natural isolate (*Vibrio campbellii*), *E. berryi* tank water | This study |
| BW2 (MJM5772) | Natural isolate (*Vibrio fischeri*), *E. berryi* tank water | This study |
| BW3 (MJM5773) | Natural isolate (*Vibrio campbellii*), *E. berryi* tank water | This study |
| BW4 (MJM5774) | Natural isolate (*Vibrio atypicus*), *E. berryi* tank water | This study |
| BW5 (MJM5775) | Natural isolate (*Vibrio campbellii*), *E. berryi* tank water | This study |
| BW6 (MJM5776) | Natural isolate (*Vibrio campbellii*), *E. berryi* tank water | This study |
| BW7 (MJM5777) | Natural isolate (*Vibrio jasicida*), *E. berryi* tank water | This study |
| BW8 (MJM5778) | Natural isolate (*Pseudoalteromonas flavipulchra*), *E. berryi* tank water | This study |
| BW9 (MJM5779) | Natural isolate (*Vibrio fischeri*), *E. berryi* tank water | This study |
| ES401 *tssF_2* (ANS2100) | ES401 *tssF_2/vasA_2* mutant (Erm<sup>R</sup>) | (71) |
| BW2 *tssF_2* (KB100) | ErmR *tssF_2/vasA_2* mutant (Erm<sup>R</sup>) | This study |
| BW2 / pSNS119 | BW2 with *vipA_2-gfp* expression vector (Kan<sup>R</sup>) | This study |
| KB100 / pSNS119 | BW2 *tssF_2* mutant with *vipA_2-gfp* expression vector (Kan<sup>R</sup>, Erm<sup>R</sup>) | This study |
| ES401 / pVSV208 | ES401-RFP (Cam<sup>R</sup>) | (27) |
| ***E. coli* strains** |  |  |
| CC118 λpir / pEVS104 (MJM534) | *E. coli* carrying the conjugation helper plasmid for triparental matings. | (72) |
| DH5α λpir / pVSV102 (MJM542) | *E. coli* carrying the GFP-expressing plasmid | (73) |

**Table 2:** Plasmids used in this study.

| Plasmid | Description | Source |
| --- | --- | --- |
| pEVS104 | Conjugation helper plasmid (Kan<sup>R</sup>) | (72) |
| pVSV102 | Constitutive GFP (Kan<sup>R</sup>) | (73) |
| pVSV208 | Constitutive DsRed (Cam<sup>R</sup>) | (73) |
| pAS2038 | *tssF_2/vasA_2* disruption vector (Erm<sup>R</sup>) | (27) |
| pSNS119 | IPTG inducible *vipA_2-gfp* expression vector (Kan<sup>R</sup>) | (27) |

### Squid monocolonizations

Hatchling squid, either *E. scolopes* or *E. berryi*, were colonized according to our previously described protocol (44) and summarized below. Bacterial strains were grown overnight with aeration in LBS at 25°C. Cultures were diluted 1:80 in LBS and subcultured until an approximate OD_600_ of 0.3. This culture was used to inoculate 40 mL of artificial seawater containing squid hatchlings, with the inoculum normalized to approximately 5 x 10^3^ CFU/mL. Squid remained in the inoculum water for 3 h at room temperature. Squid were then washed in filter-sterilized Instant Ocean (FSIO) and transferred to individual vials containing 4 mL FSIO each. Squid were transferred to fresh FSIO at 24 hpi. At 46-48 hpi, individual squid were measured for luminescence using a Promega GloMax 20/20 luminometer before euthanasia via freezing. CFU per squid was enumerated as described, wherein euthanized squid were homogenized, dilutions were plated on LBS agar, and total light organ counts were calculated from the count and proportion of colonies that were plated (44). When assaying colonization-deficient mutants, some squid in the Δ*lux* treatment group (*n* = 9) were euthanized and frozen at 24 hpi. Mutant strain CFU counts were compared to the ES114 treatment group using a Kruskal-Wallis with Dunn’s multiple comparisons test.

For light organ dissections in Fig. 3S, the above colonization protocol was adapted. At 46-48 hpi, squid were anesthetized in FSIO containing 2% ethanol and pithed. Animals were placed ventral side up and the light organ was dissected out with minimal surrounding tissue. The light organs for individual animals were then plated and enumerated as described above.

### Light organ imaging

*E. berryi* hatchlings were colonized as described above. At 48 hpi, animals were fixed in 4% paraformaldehyde in 1X mPBS (10 mL DI water, 10 mL 16% paraformaldehyde, and 20 mL 2x mPBS [marine phosphate-buffered saline: 50 mM phosphate buffer and 0.45 M NaCl [pH 7.4]). Animals remained in fixative for 48 h at 4°C. Squid were then washed 4 times for 15 minutes each at 4°C with 1X mPBS. Mantles and funnels of fixed squid were dissected away to expose the light organs of symbiotic animals. Samples were stained overnight in the dark in a permeabilization solution containing mPBS + 0.5% Triton-X 100. An overnight staining of DAPI (5 µg/µL) and Alexa647-conjugated phalloidin (25 µg/µL) was used to stain host nuclei and host F-actin, respectively. The excess stain was removed with 3 washes of mPBS. The tissues were immersed in a mPBS + 90% glycerol solution and allowed to equilibrate overnight in the dark at 4°C. Samples were transferred to 3 mm depression well slides (United Scientific), mounted ventral side facing up in the mPBS + 90% glycerol solution, and overlaid with a No. 1.5 coverslip.

Qualitative sample images were acquired on a Nikon AXR confocal microscope, illuminated with 405-, 488- and 647-nm lasers and imaged with either a 10X 0.45 NA or 20X 0.80 NA PLAN Apochromat objective. To reconstruct crypts in 3D and determine the volume occupied by bacteria, 150 µm deep Z-stacks of one lobe of each animal (*n* = 10) were acquired. The 20X 0.80 NA PLAN Apochromat objective was used, and the laser power and gain were adjusted at different focal depths using the Z Intensity Correction function (NIS Elements; Nikon) to normalize signal from the GFP-expressing bacteria across all Z-positions. Image stacks were imported to and analyzed in arivis Pro (Zeiss), where the 3D crypts were reconstructed via threshold segmentation of the GFP channel. The resulting segmented 3D objects were used to determine crypt volume.

For imaging in Fig. S3, *E. berryi* hatchlings were colonized as described above. At 48 hpi, animals were fixed in 4% paraformaldehyde in 1X mPBS. Animals remained in fixative for 24 h at 4°C. Squid were then washed 3 times for 30 minutes at 4°C with 1X mPBS. Mantles were dissected away and confirmed to have detectable GFP on the removed tissue using Zeiss Axio Zoom.V16 large-field fluorescent stereomicroscope. Samples were stained overnight in the dark at 4°C in a permeabilization solution containing mPBS + 0.5% Triton-X 100. An overnight staining of TO-PRO-3 (1 µM) was used to stain host nuclei. Excess stain was removed by washing 3 times with 1X mPBS. Samples were stored overnight at 4°C in the dark in mPBS. Samples were transferred to Corning microscope slides, mounted ventral side facing up in Vectashield Plus antifade mounting media, and overlaid with a No. 1.5 coverslip. Qualitative sample images were acquired on a Zeiss LSM 800 confocal microscope using Airyscan, illuminated with 488- and 640-nm lasers and imaged with a 20X 0.80 M27 PLAN Apochromat objective. Approximately 70 µm deep Z-stacks of each light organ lobe of each animal (*n* = 4-10) were acquired. Zen software was used for image acquisition and processing. Representative images shown are orthogonal projections of maximum intensity.

### Bacterial strain isolation and identification

Bacterial strains were isolated from *E. berryi* using two distinct methods. In the first method, luminous *E. berryi* hatchlings raised at the Marine Biological Laboratory (MBL) were euthanized, frozen, homogenized, and plated onto LBS, providing “*berryi* hatchling” (BH) isolates. As a second approach, the tank water from a wild-caught adult *E. berryi* (“*berryi* water”; BW) was collected and serial diluted onto LBS agar. This water was expected to be enriched in expelled native symbionts but also potentially included Atlantic bacteria from flow-through source water. In both approaches, bacterial isolates were streak purified on LBS, screened for luminescence, and saved as 50% glycerol stocks.

Genomic DNA was extracted from all isolates with the Qiagen DNeasy Blood & Tissue Kit. Samples were submitted to Plasmidsaurus (Eugene, OR) for bacterial genome sequencing using Oxford Nanopore Technology. Plasmidsaurus results included taxonomic assignment using sourmash (61), though this did not always provide species-level precision. Isolate species identity was then determined using FastANI v1.34 (43) against the species suggested by sourmash and likely candidates of our choosing (Table S1). Organisms belonging to the same species typically result in >95% average nucleotide identity (ANI). All bacterial isolates were found to have an ANI >95% for the listed species.

### Specific luminescence

Bacterial cultures were incubated from glycerol stocks into LBS liquid and grown overnight with aeration at 25°C. Strains were diluted 1:2 in 150 mM NaCl SWTO in a clear-bottom black 96-well optical plate using a BioTek Synergy Neo2 plate reader (62). The plate was shaken for 10 seconds before luminescence was recorded at a gain setting of 135. The plate was again shaken for 10 seconds and OD_600_ was recorded. Luminescence and OD_600_ values were normalized to blanks. Specific luminescence was calculated as luminescence-135 divided by OD_600_. Each assay was performed in triplicate, with two biological replicates conducted.

### TUNEL counts and hemocyte trafficking

*E. berryi* were inoculated as described above for monocolonizations. At 24 hpi, animals were fixed in 4% paraformaldehyde in 1X mPBS. Animals remained in fixative for 24 h at 4°C before being washed in 1X mPBS for 30 minutes each a total of 4 times at 4°C. Treatment groups had a minimum of 13 squid (*n* = 13-15). Apoptosis was assayed using the DeadEnd Fluorometric TUNEL Assay kit (Promega) according to manufacturer instructions, as described (22).

Hemocyte counts were taken from the anterior appendage of individual *E. berryi*, adapted from the method described (24, 63). Briefly, we stained fixed animals with FITC-DnaseI overnight, counterstained with TOTO-3 to visualize nuclei, and then visualized samples on a Lecia Stellaris 5 CLSM. A non-parametric one-way ANOVA (Kruskal-Wallis) using Dunn’s multiple comparisons test was used to compare each symbiotic group to the aposymbiotic control group.

### T6SS2 Assays

The BW2 *tssF* mutant was made by introducing a disruption into *tssF_2* in BW2 through triparental mating and *E. coli* carrying the *tssF_2* disruption plasmid pAS2038 (27). Because the *tssF_2* gene, which encodes an essential structural protein in the T6SS gene cluster on chromosome II (T6SS2), is conserved across different strains, but distinct from the *tssF* gene encoded in the T6SS cluster on chromosome I (T6SS1), we can use the same *tssF_2* disruption plasmid (pAS2038) to generate a T6SS2 mutant in diverse strains while leaving T6SS1 intact. Fluorescently-tagged strains were made by triparental mating, introducing pVSV102 for GFP-tagged variants and pVSV208 for dsRed-tagged variants. VipA-GFP expressing strains were made by introducing pSNS119 (27) using triparental mating.

We previously identified a high-viscosity condition that mimics the host environment and enables visualization of T6SS sheaths *in vitro* (64). To visualize T6SS sheath formation, strains carrying the IPTG-inducible VipA_2-GFP expression vector (pSNS119) were grown in LBS liquid supplemented with kanamycin overnight with shaking at 24°C. Cultures were then diluted 1:80 into fresh LBS supplemented with 5% w/v polyvinylpyrrolidone (PVP) and 0.5 mM isopropyl-β-d-thiogalactopyranoside (IPTG) and grown to an OD_600_ of 1.0 (64). Two microliters of each culture was spotted onto a 35 mm petri dish with #1.5 glass bottom, and a 2% agarose pad with 0.5 mM IPTG was placed on top of the culture. Cells were imaged using a Nikon Ti2 inverted fluorescence microscope outfitted with a Hamamatsu Orca Fusion C1440 and CFI plan apo lambda 100X oil objective lens. Images were captured using NIS-element software and brightness and contrast was adjusted using FIJI.

Co-incubation assays were performed between the indicated strain and GFP-tagged ES114 as described (65).

Competitive colonizations were performed as described for single-strain monocolonizations, except both designated strains were provided as inoculum. Strains were differentiated for colony counting by fluorescent label, using ES114-GFP and ES401-RFP, or the respective ES401 derivative with RFP.

### Data analysis

Plotting and statistical analysis were conducted in GraphPad Prism. All data analyses were performed using nonparametric tests. Normality tests in Prism (Kolmogorov-Smirnov, Shapiro-Wilk, and Anderson-Darling tests) indicated for all datasets that data were not normally distributed or the sample size was too small to determine normality. Affinity designer was used for figure design.

Cam, chloramphenicol; Erm, erythromycin; Kan, kanamycin; GFP, green fluorescent protein; DsRed, Discosoma sp. red fluorescent protein; IPTG, isopropyl-β-d-thiogalactopyranoside.

## Supplemental Material

**Supplemental Table S1:** Reference genomes used in FastANI analysis for species identification.

**Supplemental Figure S1.** Comparison of adult mantle length.

**Supplemental Figure S2.** ES114 occupies a greater crypt volume in crypt 1 than BW2.

**Supplemental Figure S3.** Non-*V*. fischeri are not observed in the *E. berryi* light organ or surrounding tissues.

**Supplemental Figure S4.** Inoculum concentration did not positively correlate to squid colonization concentration in strains that exhibited bimodality.

**Supplemental Figure S5.** *E*. scolopes are colonized by *V. fischeri* isolated from *E. berryi*.

## ACKNOWLEDGMENTS

We are grateful to Joshua Rosenthal for assistance throughout this project; to Denise Ludvik for support with imaging; to Ruth Isenberg for construction of MJM3972; and to Kewalo Marine Laboratory for *Euprymna scolopes* field resources. Work in the lab of MJM was supported by NIGMS R35GM148385, in the lab of ANS by NIGMS R35GM137886, in the lab of EACH by NIGMS R35GM150478, and in the lab of SVN by NSF IOS-2247195 and the Gordon and Betty Moore Foundation. AMI was supported on NIGMS T32GM007133.

