## Supplemental Material for "*Euprymna berryi* as a comparative model host for *Vibrio fischeri* light organ symbiosis"

**Supplemental Table S1:** Reference genomes used in FastANI analysis for species identification.

| Species | Accession number |
|---|---|
| *Pseudoalteromonas flavipulchra* | ASM25911v1 |
| *Pseudoalteromonas maricaloris* | ASM2474685v1 |
| *Vibrio atypicus* (DSM 25292) | ASM981131v1 |
| *Vibrio campbellii* | ASM290647v1 |
| *Vibrio fischeri* (ES114) | ASM1180v1 |
| *Vibrio fischeri* (MB14A3) | ASM164037v1 |
| *Vibrio fischeri* (MJ11) | ASM2084v1 |
| *Vibrio harveyi* | ASM3006043v1 |
| *Vibrio parahaemolyticus* | ASM19609v1 |
| *Vibrio vulnificus* | ASM222426v1 |

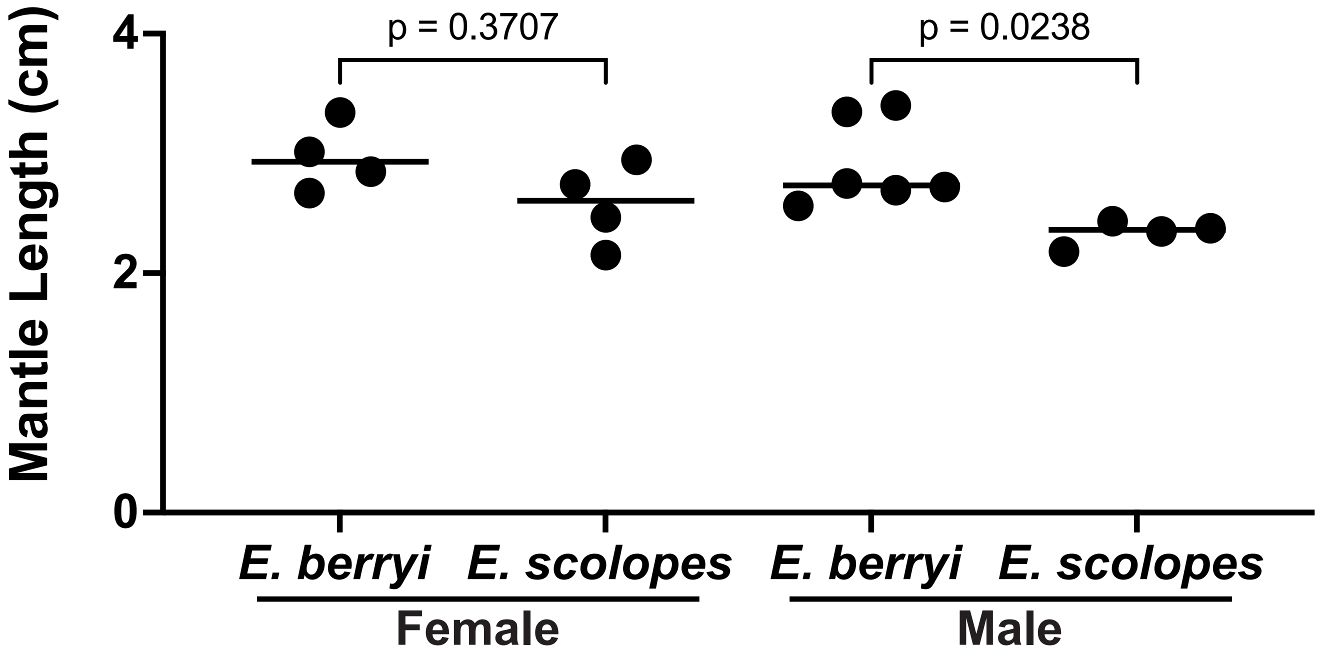


**Supplemental Figure S1. Comparison of adult mantle length.** Mantle lengths of adult *Euprymna* held in laboratory aquaculture. Mantle lengths were estimated in FIJI. Circles represent individual animals and the median is plotted ($n$ = 4-6). A Kruskal-Wallis test was applied for statistical analysis. Among our small sample, we observed a statistically significant difference among males a trend among females where *E. berryi* was larger.

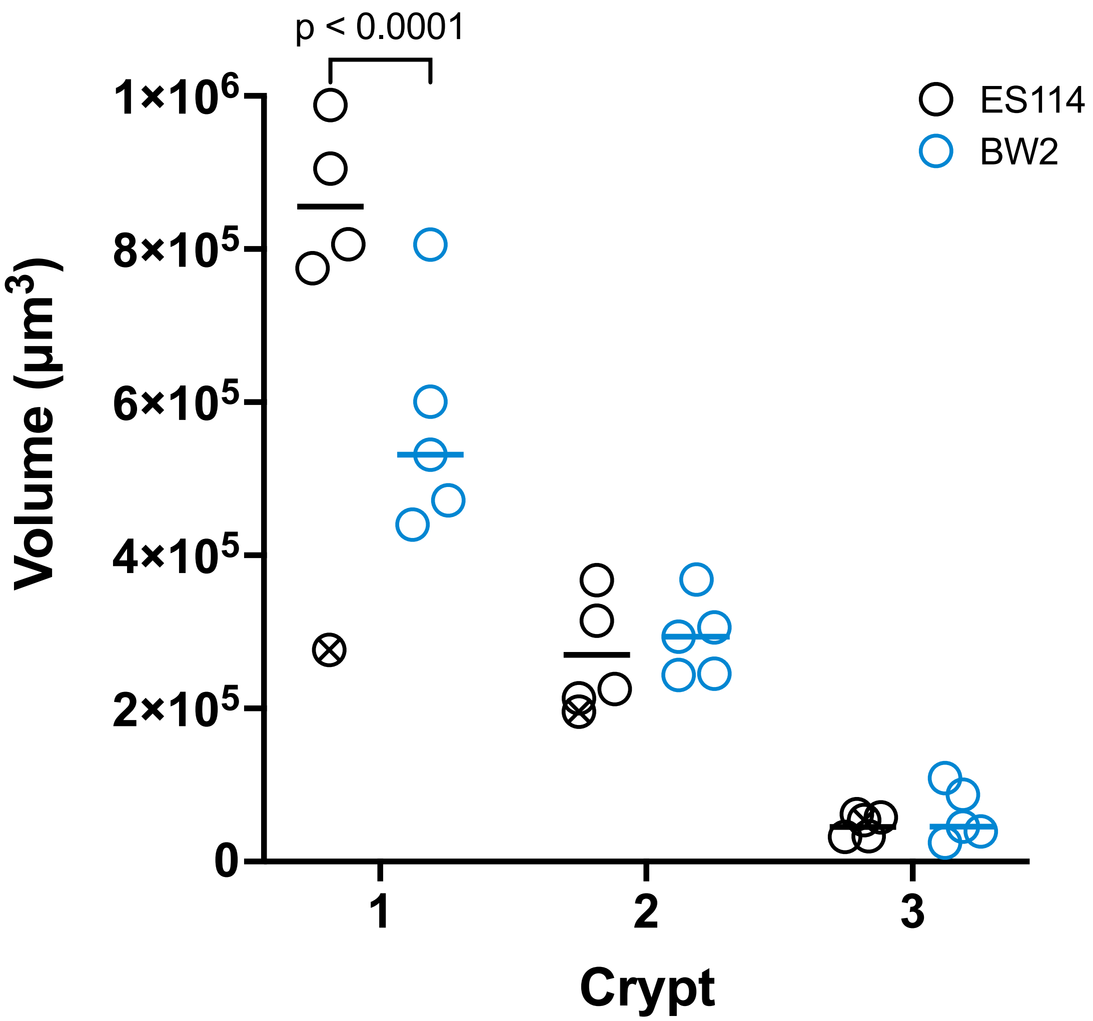


**Supplemental Figure S2. ES114 occupies a greater crypt volume in crypt 1 than BW2.** Z-stack images of single light organ lobes of *E. berryi* approximately 150 µm deep were taken at 1.2 µm intervals. Animals were colonized with a pVSV102 GFP-expressing derivative of the indicated strain. ES114 is a *V. fischeri* isolate from *E. scolopes,* and BW2 is a *V. fischeri* isolate from *E. berryi*. Image stacks were processed and analyzed using Arivis software, using the GFP signal to determine the space occupied by the *V. fischeri* and calculate the size of the crypt. The median of each group is shown with a horizontal bar (*n* = 4-5). ES8, an animal that vented its crypt bacteria during processing, is marked with an X and omitted from statistical analysis. Statistical analysis was performed using a two-way ANOVA test. Crypts 2 and 3 were not significantly different ($p > 0.99$).

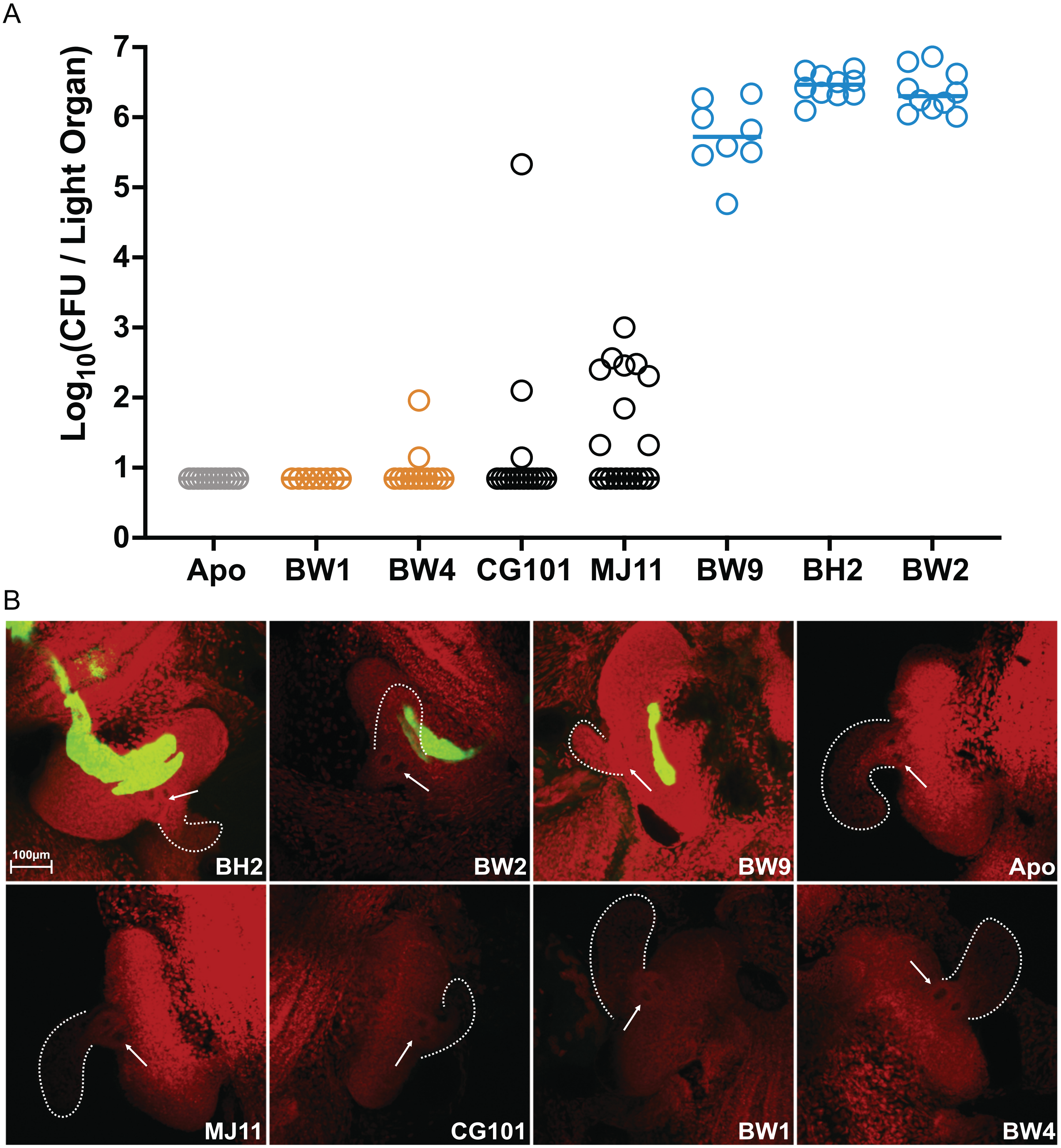


**Supplemental Figure S3. Non-*V. fischeri* are not observed in the *E. berryi* light organ or surrounding tissues.** (A) *E. berryi* hatchlings were used in single-strain colonization experiments, where circles represent individual animals. Colonization was estimated by dissection of the light organ and minimal surrounding tissue for homogenization and plating. The limit of detection for this assay, represented by the dashed line, is 7 CFU/light organ. Horizontal bars represent the median for each group ($n$ = 12-19). (B) Representative images of *E. berryi* inoculated with the indicated GFP-expressing (green) bacterial strain ($n$ = 4-10). Images are a maximum intensity projection of a confocal z-stack approximately 70 µm deep. Host tissue is stained with TO-PRO-3 (red). The anterior appendage is outlined in a dotted white line. The three pores of each lobe are indicated by a white arrow. All images have the same scale.

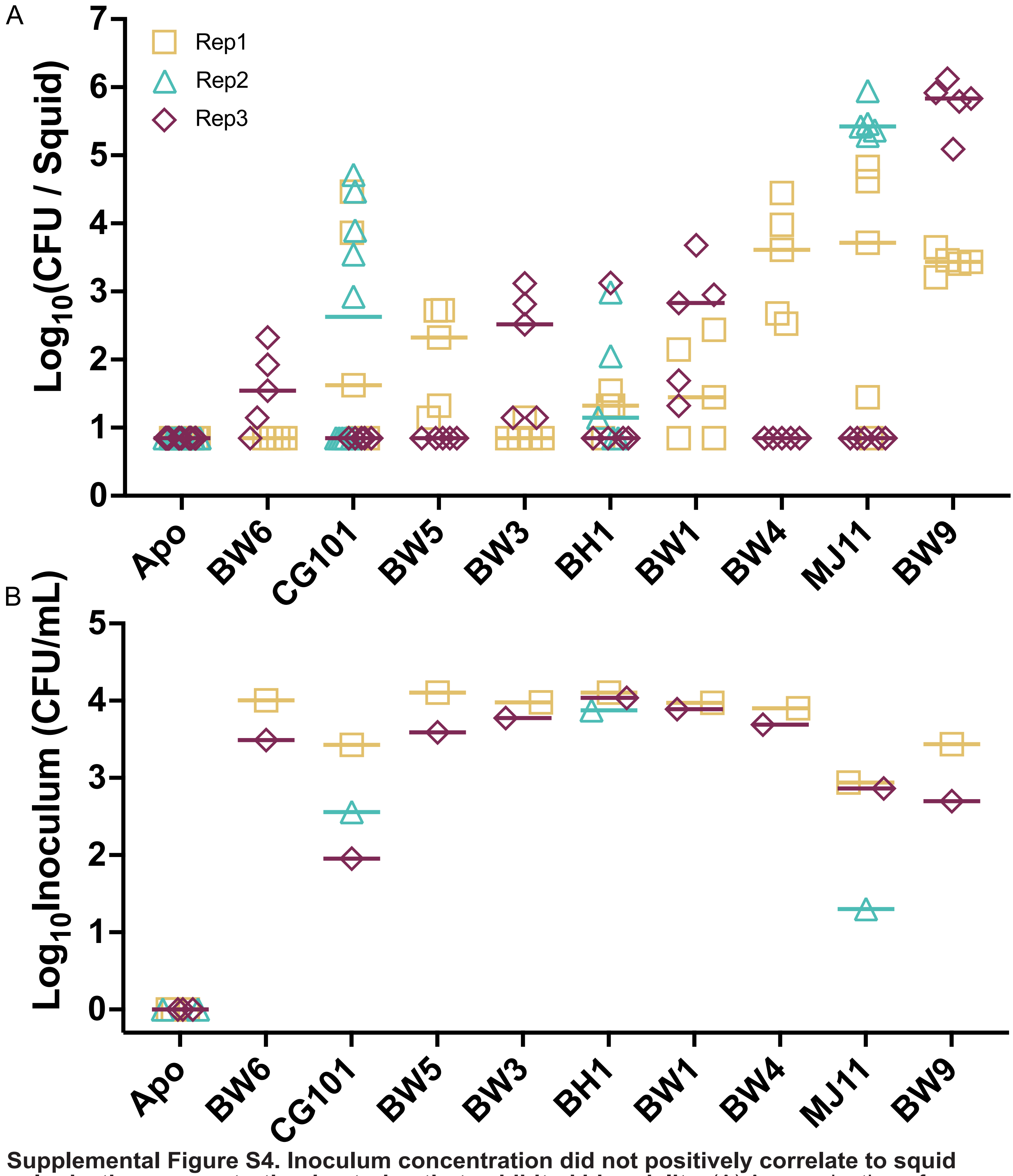


**Supplemental Figure S4. Inoculum concentration did not positively correlate to squid colonization concentration in strains that exhibited bimodality.** (A) A reproduction of monocolonization data for select strains as presented in Fig4. Each symbol represents an individual animal, where color and shape indicate the replicate group that animal belonged to. Apo represents aposymbiotic squid, which are not provided bacterial inoculum in these experiments. (B) The inoculum concentration provided to squid, where each symbol represents the concentration of the indicated strain provided to the squid group within an experiment. Horizontal bars represent the median in both graphs. Colors in panel A and B correspond to the same replicate across experiments.

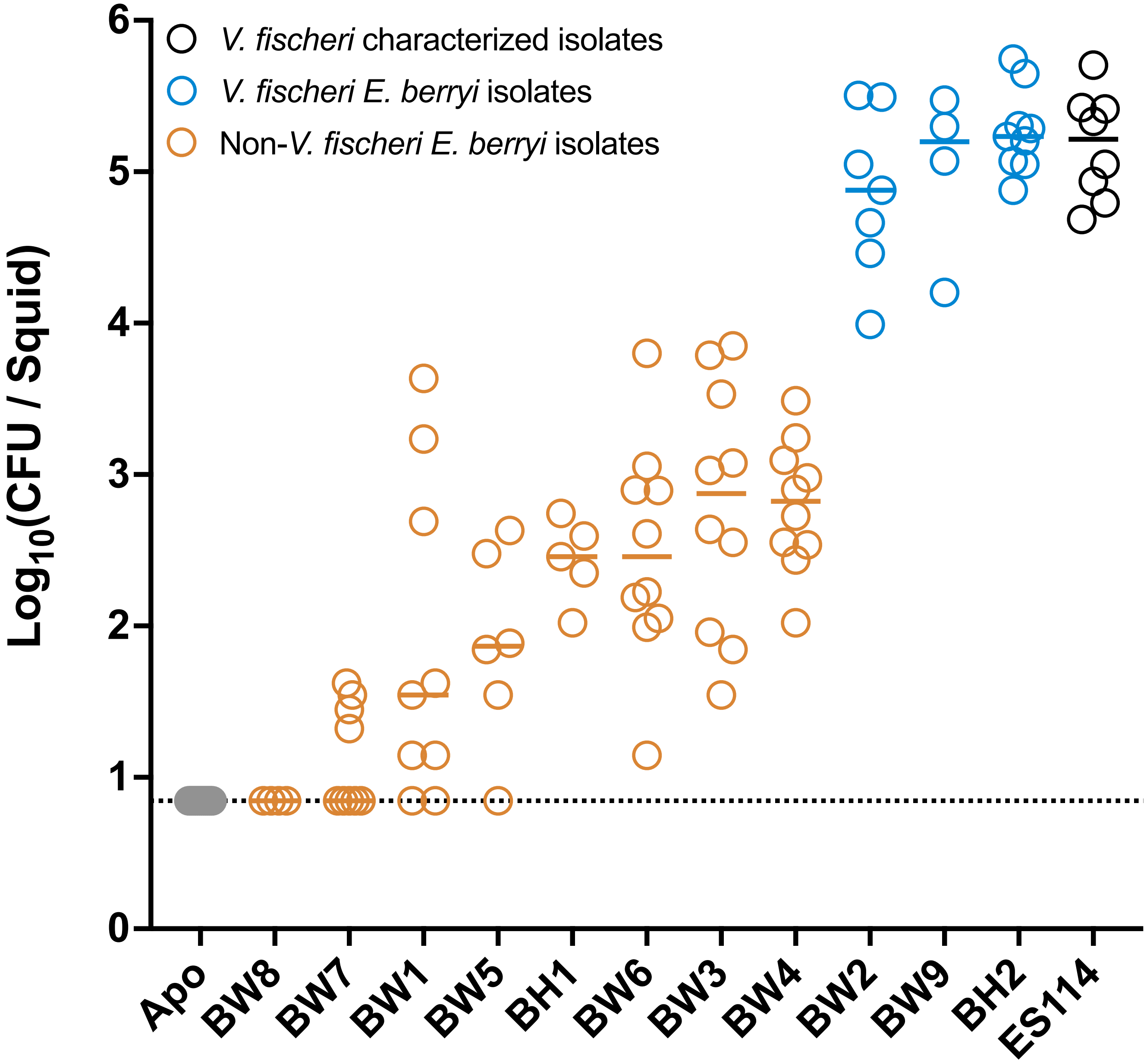


**Supplemental Figure S5. *E. scolopes* are colonized by *V. fischeri* isolated from *E. berryi*.** *E. scolopes* hatchlings were used in single-strain colonization experiments, where circles represent individual animals. The limit of detection for this assay, represented by the dashed line, is 7 CFU/light organ. Horizontal bars represent the median for each group (*n* = 4-13).
